# Ca^2+^ Mediates HIF-dependent Upregulation of Aquaporin 1 in Pulmonary Arterial Smooth Muscle Cells

**DOI:** 10.1101/2021.12.06.471473

**Authors:** Xin Yun, Stephan Maman, Haiyang Jiang, Gregg L. Semenza, Larissa A. Shimoda

## Abstract

Prolonged exposure to hypoxia causes structural remodeling and sustained contraction of the pulmonary vasculature, resulting in the development of pulmonary hypertension. Both pulmonary arterial smooth muscle cell (PASMC) proliferation and migration contribute to the vascular remodeling. We previously showed that the protein expression of aquaporin 1 (AQP1), a membrane water channel protein, is elevated in PASMCs during following in vivo or in vitro exposure to hypoxia. Studies in other cell types suggest that AQP1 is a direct transcriptional target of hypoxia inducible factor (HIF)-1. Moreover, we and others have shown that an increase in intracellular calcium concentration ([Ca^2+^]_i_) is a hallmark of hypoxic exposure in PASMCs. Thus, we wanted to determine whether HIF regulates AQP1 in PASMCs and, if so, whether the process occurred via transcriptional regulation or was Ca^2+^-dependent. PASMCs were exposed to hypoxia, incubated with DMOG, which inhibits HIFα protein degradation or infected with constitutively active forms of HIF-1α or HIF-2α. Hypoxia, DMOG and HIF1/2α produced a time-dependent increase in AQP1 protein, but not mRNA. Interestingly, incubation with increasing HIF1/2α levels and DMOG increased [Ca^2+^]_i_ in PASMCs, and this elevation was prevented by the voltage-gated Ca^2+^ channel inhibitor, verapamil (VER) and nonselective cation channel inhibitor SKF96365 (SKF). VER and SKF also blocked upregulation of AQP1 protein by DMOG or HIF1/2α, but had no effect on expression of GLUT1, a canonical HIF transcriptional target. Silencing of AQP1 abrogated increases in PASMC migration and proliferation induced by HIF1/2α, suggesting induction of AQP1 protein by HIF1/2α has a functional outcome in these cells. Thus, our results show that contrary to reports in other cell types, in PASMCs, AQP1 does not appear to be a direct target for HIF transcriptional regulation. Instead, AQP1 protein may be upregulated by a mechanism involving HIF-dependent increases in [Ca^2+^]_i_.

## INTRODUCTION

Pulmonary hypertension (PH), defined as pulmonary arterial pressure > 20 mmHg (30), is a complex condition arising from various etiologies, including genetic and environmental factors. For example, PH is a common consequence of exposure to chronic hypoxia due to residence at high altitude, sleep apnea or chronic lung diseases such as emphysema or chronic bronchitis. Remodeling of the pulmonary vascular wall and contraction of the pulmonary arteries contribute to the elevations in pulmonary arterial pressure regardless of inciting cause. With chronic hypoxia, both of these processes are associated with enhanced proliferation and migration of pulmonary arterial smooth muscle cells (PASMCs) which results in thickening of the medial layer and appearance of smooth muscle in normally non-muscular arterioles (26). While these structural and functional abnormalities in the pulmonary hypertensive vasculature have been well documented for decades, the underlying cellular mechanisms are complicated and incompletely understood.

Recent work has established a role for aquaporin 1 (AQP1) in mediating PH induced by exposure to chronic hypoxia (15, 16, 18, 25, 40). Aquaporins (AQPs) comprise a family of transmembrane proteins found throughout the body that function as water channels, with AQP1 being the first discovered (41). In addition to a canonical role in transporting water across membranes, AQP1 also regulates the accumulation of β-catenin in mesenchymal stem cells, breast cancer, endothelial and melanoma cells (7, 21, 22). We documented a similar AQP1-induced increase in β-catenin in PASMCs, a process we found to be independent of water transport but requires the AQP1 C-terminal tail region (40). Interestingly, increased abundance of AQP1 is sufficient to induce changes in PASMC phenotype (15, 25), suggesting that a better understanding of the mechanisms by which AQP1 protein levels are regulated could be important for development of therapeutics to target vascular remodeling. While several labs, including our own, showed that AQP1 protein is upregulated in the smooth muscle cells of rodents exposed to chronic hypoxia and contributes to enhanced proliferation and migration (16, 18, 25), the exact mechanism by which hypoxia induces AQP1 expression in PASMCs has not been established.

Several studies demonstrated a role for hypoxia inducible factors (HIFs) in direct transcriptional regulation of AQP1 in retinal endothelial and hemangioendothelioma cells (1, 31). In our previous work in PASMCs, while AQP1 mRNA levels were increased in PASMCs isolated from the chronic hypoxia model, we showed that the initial upregulation of AQP1 in response to in vitro exposure to hypoxia occurred without an increase in mRNA levels(16), suggesting the potential for a different post-transcriptional regulatory mechanism in these cells. Thus, whether the upregulation of AQP1 protein is a consequence of HIF activation and mediated via transcriptional regulation in smooth muscle remains uncertain. In the current study, our goal was to determine whether the hypoxia-induced upregulation of AQP1 in PASMCs is HIF-dependent and, if so, the downstream mechanism(s) involved.

## METHODS

All protocols were reviewed by and performed in accordance with Johns Hopkins University Animal Care and Use Committee. Protocols and procedures comply with NIH and Johns Hopkins Guidelines for the care and use of laboratory animals.

### Exposure of Mice to Chronic Hypoxia

In this study, we performed measurements in samples collected and frozen in a previously reported study (2). Briefly, adult male C57B/6J mice (8-10 weeks, Jackson Labs) were randomly assigned to one of 4 groups: normoxia+vehicle, normoxia +digoxin, chronic hypoxia+vehicle or chronic hypoxia +digoxin. Mice were exposed to hypoxia in a chamber maintained at 10% O2 for 3 wk. The chamber was constantly flushed with room air to maintain low (<0.5%) CO_2_ concentrations. A servo-control system (PRO-OX; Hudson RCI) monitored O_2_ levels and injected 100% N_2_ as needed to maintain 10% ± 0.5% O_2_. Cages were cleaned and food and water replenished twice per week. Normoxic animals were kept in room air on a wire rack adjacent to the chamber. All animals were allowed free access to food and water. Beginning one day prior to hypoxic exposure, mice were weighed and injected daily with digoxin (1.0 mg/kg i.p.; diluted in sterile saline) or vehicle (sterile saline). Injectable digoxin (Sandoz; 0.25 mg/mL) was obtained from The Johns Hopkins Hospital Research Pharmacy. At the end of exposure, lung tissue samples were harvested by an investigator blinded to treatment groups and snap frozen in liquid nitrogen and stored at −80°C until use.

### Isolation and culture of PASMCs

Adult Wistar rats (250-300g) were obtained from Harlan Farms and were anesthetized with pentobarbital sodium (130 mg/kg i.p.). Because female rats are less susceptible to the effects of hypoxia, only male rats were used in this study. Under deep anesthesia, the chest was opened and the heart and lungs excised and placed in a dissection dish containing ice cold HEPES-buffered saline solution (HBSS) (130 mM NaCl, 5 mM KCl, 1.2 mM MgCl_2_, 1.5 mM CaCl_2_, 10 mM HEPES and 10 mM glucose, with pH adjusted to 7.2 using 5 M NaOH. Resistance level pulmonary arteries (200-600 μM outer diameter) were isolated, the adventitia removed and opened for endothelial denudation with a cotton-topped swab. Tissue was placed in 4°C HBSS for at least 30 min, transferred to room temperature reduced-Ca^2+^ HBSS (20 μm CaCl_2_) for at least 20 min and then digested in reduced-Ca^2+^ HBSS containing: type I collagenase (1750 U/ml), papain (9.5 U/ml), bovine serum albumin (2 mg/ml), and DTT (1 mM) at 37°C for 15-20 min. After digestion, the tissue was transferred to Ca^2+^-free HBSS in a small round bottom tube and slowly triturated to create a single cell suspension. Dispersed PASMCs were cultured in basal SmBM media (Lonza) supplemented with 0.3% FCS and 1% penicillin-streptomycin for 48 h, followed by 3-4 days of culture in SmGM complete media (Lonza) supplemented with 1% penicillin-streptomycin to expand cell numbers. PASMCs were starved in SmBM media for 24-48 h before beginning experiments. All cells were used at passage 0. Assessment of smooth muscle culture purity was performed using: 1) [Ca^2+^]_i_ responses to 80 mM KCl and 2) immunofluorescence in cells stained with smooth muscle-specific α-actin (SMA; 1:400, A2547, Sigma), calponin (1:100, ab700, Abcam) or smooth muscle myosin heavy chain (SMMHC; 1:100, ab125884, Abcam) and DAPI (nuclear stain; 1:10,000 in PBS, Invitrogen). Only cultures where >90% of cells exhibited at least 50 nM increase in [Ca^2+^]_i_ and positive SMA and calponin or SMMHC expression were used for these studies.

### In vitro hypoxic exposure

Exposure of PASMCs to hypoxia was achieved by placing tissue culture plates containing cells in a modular chamber (Billups-Rothberg) gassed with 4% O_2_; 5% CO_2_ as previously described (16). The chamber containing cells and the normoxic control plates were placed in a room air incubator maintained at 37°C and 5% CO_2_. Correct oxygen levels inside the hypoxic chamber were confirmed using a hand-held oxygen monitor (model 5577; Hudson RCI).

### Real-time RT-PCR

RNA was extracted from PASMCs using the RNeasy Plus Mini kit (Qiagen) following manufacture’s instructions. Reverse transcription was performed using 500 ng of RNA and the iScript cDNA synthesis kit (Bio-Rad). mRNA expression levels were measured using real-time PCR, which was performed using 500 ng of cDNA with QuantiTect SYBR Green PCR Master Mix (Qiagen) in an iCycler IQ real-time PCR detection system. The specific primer pairs for real-time PCR are listed in **Table 1**. PCR products were confirmed by: 1) a single peak in the melt curve; 2) a single band of the correct size when products were run on an agarose gel and 3) sequencing of the product. Relative concentrations of each gene were calculated using the Pfaffl method, and data are expressed as a power ratio of the gene of interest to the housekeeping gene (Cyclophilin B; CpB) within a sample using the efficiency for each gene calculated from standard curves run on the same plate as samples. For each experiment, the values were normalized to the value of control sample.

**Table 1:**
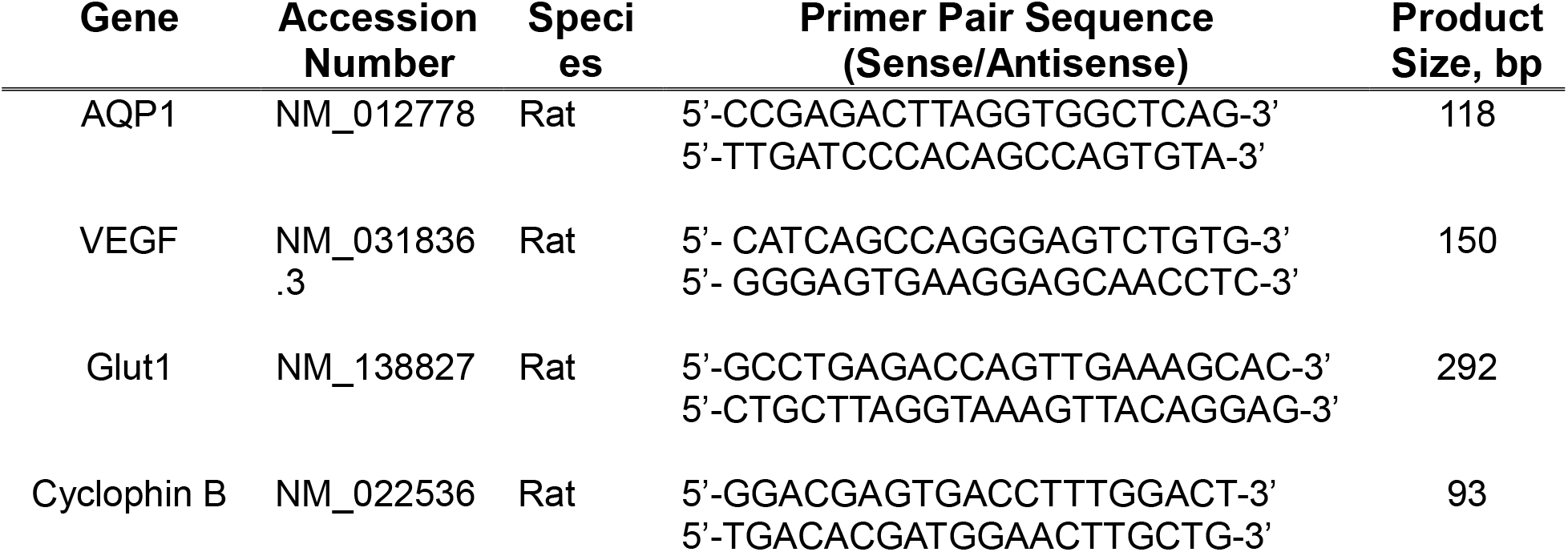
Primers for PCR.

### Western Blotting

PASMCs were washed with PBS, and total protein extracted in ice-cold T-PER buffer containing protease inhibitors (Roche Diagnostics). Proteins were quantified by use of the BCA protein assay (Pierce) and 8-20 μg total protein were resolved by 10% SDS-PAGE gels transferred onto polyvinylidene difluoride membranes, which were then blocked with 5% non-fat dry milk in Trisbuffered saline containing 0.2% Tween 20. Membranes were probed with primary antibodies AQP1 at 1:4,000 (AQP11A, Alpha Diagnostic Intl. Inc.), glucose transporter 1 (GLUT1) at 1:1,000 (ab115730, Abcam), HIF-1α (1:1000, ab18471, Abcam) or HIF-2α (1:1000, ab8365, Abcam). Antibodies are verified with siRNA knockout by our lab (AQP1) previously (15) or the antibody company (GLUT1), or by viral overexpression in this manuscript (HIF-1α and HIF-2α). Bound antibodies were probed with horseradish peroxidase-conjugated anti-rabbit or anti-mouse IgG (1:10,000, 52200336 and 52200341, Kirkegaard & Perry Laboratories) and detected by enhanced chemiluminescence. Membranes were then stripped and re-probed for β-tubulin (1:10,000, T7816, Sigma) as a housekeeping protein. Protein levels were quantified by densitometry using ImageJ.

### Intracellular Ca^2+^ Measurements

PASMCs were placed in a laminar flow cell chamber perfused with modified Kreb’s solution (KRB) containing (in mM): 118.3 NaCl, 4.7 KCl, 1.2 MgSO_4_, 25 NaHCO_3_, 11 glucose and 1.2 KH2PO4, and gassed with 16% O2; 5% CO2 or with HBSS with pH adjusted to 7.4 as described previously (2, 28). Cells were incubated with the membrane permeant (acetoxymethyl ester) form of the Ca^2+^-sensitive fluorescent dye Fura-2 AM for 60 min at 37° C under an atmosphere of 21% O_2_; 5% CO_2_. Cells were then washed with KRB for 15 min at 37° C to remove extracellular dye and allow complete de-esterification of cytosolic dye. Ratiometric measurement of Fura-2 fluorescence was performed on a workstation (Intracellular Imaging Inc, Cincinnati, OH) consisting of a Nikon TSE 100 Ellipse inverted microscope with epi-fluorescence attachments. Light from a xenon arc lamp was alternately filtered by 340 nm and 380 nm interference filters and focused onto PASMCs via a 20x fluorescence objective (Super Fluor 20, Nikon). A filter cube was used to collect light emitted from the cell at 510 nm. Filtered light was then returned through the objective and detected by a cooled CCD imaging camera. Between measurements, an electronic shutter (Sutter Instruments) was used to minimize photobleaching of eye. All protocols were performed and data collected on-line with InCyte software (Intracellular Imaging Inc, Cincinnati, OH). The ratio of 340 to 380 nm emission was calculated by the software and [Ca^2+^]_i_ was estimated using an *in vitro* calibration curve.

### Adenovirus Infection

Adenoviral constructs containing a hemagglutinin (HA)-tagged constitutively active form of HIF-1α (NM_001530.3; P402A, P564A) or HIF-2α (NM_001430.4; P405A, P531A) were obtained from Vectorbuilder (Chicago, IL). The sequences of the plasmids encoding these vectors were verified by Sanger sequencing (The Genetics Resources Core Facility, Johns Hopkins University). The titer of the virus was determined by Adeno-X Rapid Titer Kit (Clontech). Cells were infected with 50 MOI for 48 hr. The same adenoviral vector containing EGFP (AdEGPF) was used as a control.

### siRNA

Depletion of endogenous AQP1 was achieved using siRNA targeting specific proteins and non-targeting siRNA (siNT; control) obtained as a “smart pool” (Dharmacon) as previously described (16). PASMCs were incubated with 100 nM of siRNA for 16 h in serum- and antibiotic-free media, after which serum was added to media for a total concentration of 0.3% FCS. Cells were incubated under these conditions for 24 h, and then media was replaced and cells were incubated for an additional 24 h in basal media (0.3% FCS) prior to experiments.

### Drugs

The HIF-1 inhibitor, CAY10585 (Cayman), was made as a 60 mM stock solution in DMSO. The HIF-2 inhibitor, TC-S 7009 (Tocris), was made as a 20 mM stock solution in DMSO. Both verapamil (VER; V4629) and SKF-96365 (SKF; S7809) were obtained from Sigma-Aldrich and made as 10 mM stock solutions. All stock solutions were aliquoted and frozen for storage, and diluted to working concentrations in perfusate or medium on the day of experiment.

### Cell Migration

50,000 cells in low serum media (0.3% FCS) were added to the top chamber of a polycarbonate transwell insert (8 μm pores) inserted into 6-well plates. Cells were allowed to migrate in low serum media for 24 h, after which the cells were fixed in 95% ethanol for 10 min, stained with Brilliant Blue (Pierce) for 5 min, and visualized via a microscope-mounted camera. For each filter, five fields were randomly chosen and imaged with Q-capture software. Unmigrated cells were then scraped off from the top of the filter and the bottom layer re-imaged (migrated cells). Migration rate was calculated as the percent of cells remaining on the filter after scraping normalized to the total amount of cells.

### Cell Proliferation Assay

5,000 cells were seeded into low-serum media (0.3% FCS) in wells of a 96-well plate in triplicate for every sample. After incubating for 2-3 h to allow cells to adhere, BrdU (Amersham Biosciences) was added to each well for 24 h, after which the media was removed and cells were fixed. Proliferation was estimated from detection of peroxidase-labelled anti-BrdU in newly synthesized cells. Developed color was measured at 450 nm in a microtitre plate spectrophotometer. Proliferation rate is shown as percent of the absorbance values normalized to the control cells on the same plate, with control values set to 100%.

### Statistical analysis

Data are expressed as scatter plots with bars representing means ± SD. Each dot represents a separate experimental run, and since all experimental runs were performed on tissue/cells from different animals, “n” also refers to the number of animals. For [Ca^2+^]_i_ experiments, data was collected from 10-30 cells per coverslip, and averaged to obtain a single biological replicate per experiment. All data were tested for normality prior to running statistical tests. Statistical comparisons were performed using Students t-test for data in two groups, or one- or two-way ANOVA with a Holm-Sidak post hoc test for multiple group comparisons.

## RESULTS

### Role of HIFs in hypoxia-induced upregulation of AQP1 protein expression

In a previous study, we demonstrated treatment of mice with digoxin, to disrupt HIF-1α accumulation (42), prevented the development of hypoxia-induced pulmonary hypertension (2). To determine whether AQP1 protein levels were increased in the lungs of chronically hypoxic mice and, if so, whether this increase was attenuated by treatment with digoxin, we assessed protein levels of AQP1 in lung samples collected during the prior experiment. Treatment with digoxin had no effect on baseline AQP1 protein levels in normoxic animals (**Fig 1A**). AQP1 levels were elevated in vehicle-treated mice exposed to chronic hypoxia, but not in chronically mice treated with digoxin. These results are consistent with our previous findings in rats demonstrating that exposure to chronic hypoxia increases AQP1 levels in the lung (16), and further suggests a role for HIFs in mediating the response.

**Figure 1.**
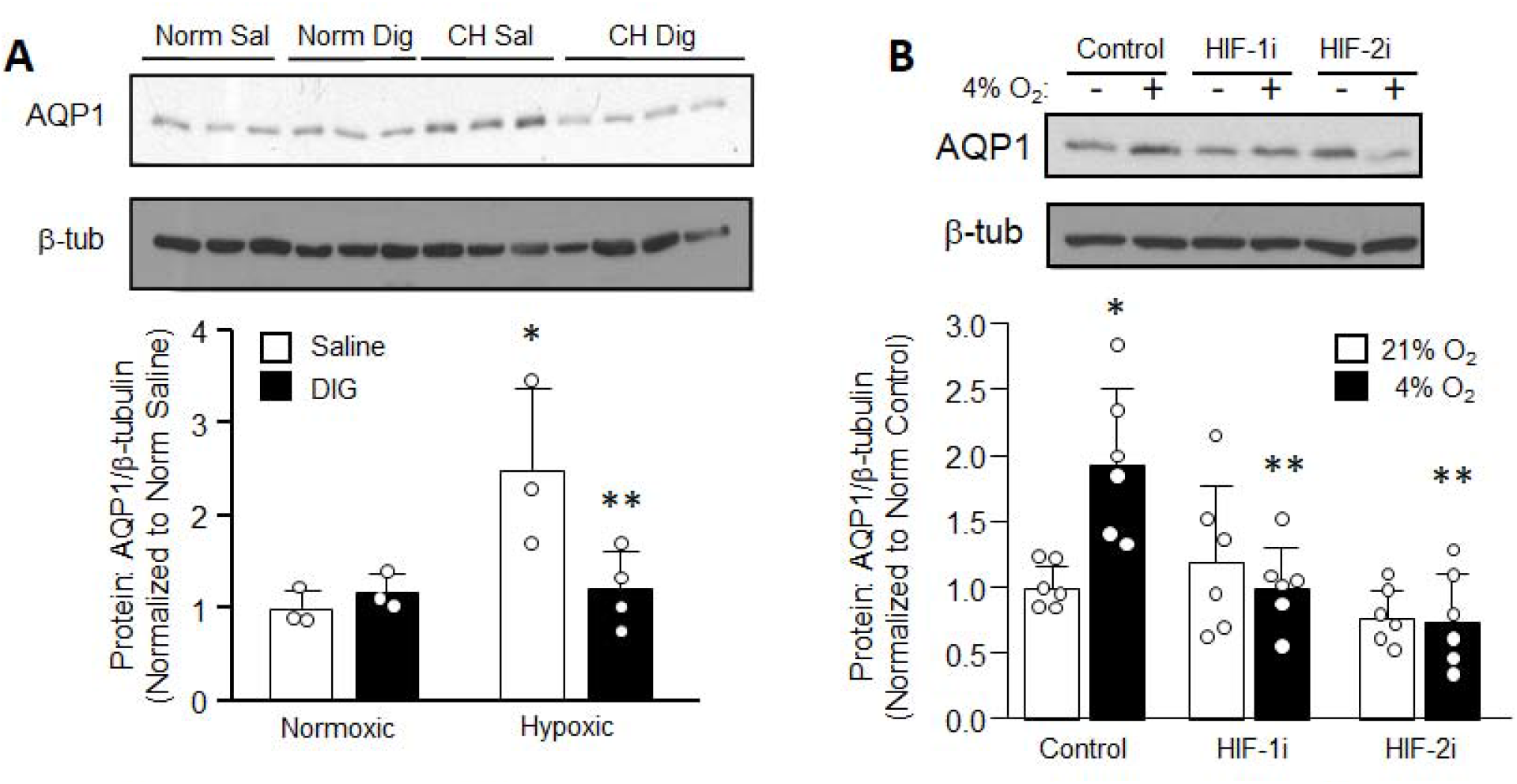
Hypoxia-inducible factor (HIF)-dependent upregulation of aquaporin 1 (AQP1) during hypoxia A) Representative immunoblot and bar (mean ± SD) and scatter plots showing AQP1 protein levels in lung tissue from normoxic (Norm) and chronically hypoxic (CH) mice treated with saline (Sal) or digoxin (0.2 mg/kg/day). p=0.029 for interaction by 2-way ANOVA. * indicates p<0.05 from Norm Sal; ** indicates p<0.05 from CH Sal by Holm-Sidak post-test B) Representative immunoblot and bar (mean ± SD) and scatter plots showing AQP1 protein expression in pulmonary arterial smooth muscle cells from control rats exposed to control (21% O_2_) or hypoxic (4% O_2_) conditions for 48 h in the presence of vehicle (1:2000 DMSO) or inhibitors for HIF-1 (HIF-1 i; 30 μM CAY10585) or HIF-2 (HIF-2i; 10 μM TC-S 7009). * indicates p<0.001 from Control 21% O_2_; “ indicates p<0.001 from Control 4% O_2_ via one-way ANOVA with Holm-Sidak post-test. Each dot represents a biological replicate using tissue/cells isolated from a different animal.

To confirm a role for HIFs in mediating the effect of hypoxia on AQP1 levels, we next challenged rat PASMCs with in vitro hypoxic exposure in the absence and presence of specific small molecular inhibitors for HIF-1 (CAY10585; 30 μM) and HIF-2 (TC-S 7009; 10 μM). Consistent with our previous results (16), short-term (48 h) in vitro exposure to hypoxia increased AQP1 protein levels in PASMCs (**Fig 1B**). Treatment with inhibitors for either HIF-1 or HIF-2 completely prevented the hypoxia-induced increase in AQP1 protein. In some experiments, HIF inhibitors also appeared to reduce normoxic levels of protein, but this was not a consistent finding across all experiments (p=0.78 for HIF-1 inhibitor and 0.93 for HIF-2 inhibitor).

### Effect of increasing HIF activity on AQP1 expression in PASMCs

We next tested whether inhibiting prolyl hydroxylases (PHDs) with DMOG, which increases HIF-1α and HIF-2α accumulation, was sufficient to mimic hypoxia and increase AQP1 levels in PASMCs. Using immunoblot analysis, we found that DMOG treatment for 24 h had a variable effect on AQP1 protein levels (p=0.841; **Fig 2A**). However, prolonging the treatment time to 48 hr caused a consistent increase in AQP1 protein levels. Interestingly, in the same cells, DMOG markedly upregulated the protein levels of GLUT1 (**Fig 2B**), a canonical HIF transcriptional target, at both 24 and 48 hr. The effect of DMOG on AQP1 expression was restricted to protein levels, as no consistent effect on AQP1 mRNA levels was evident in PASMCs at either 24 or 48 hr (**Fig. 2C**). In contrast, DMOG caused a striking and expected upregulation of GLUT1 mRNA levels at both time points (**Fig 2D**).

**Figure 2.**
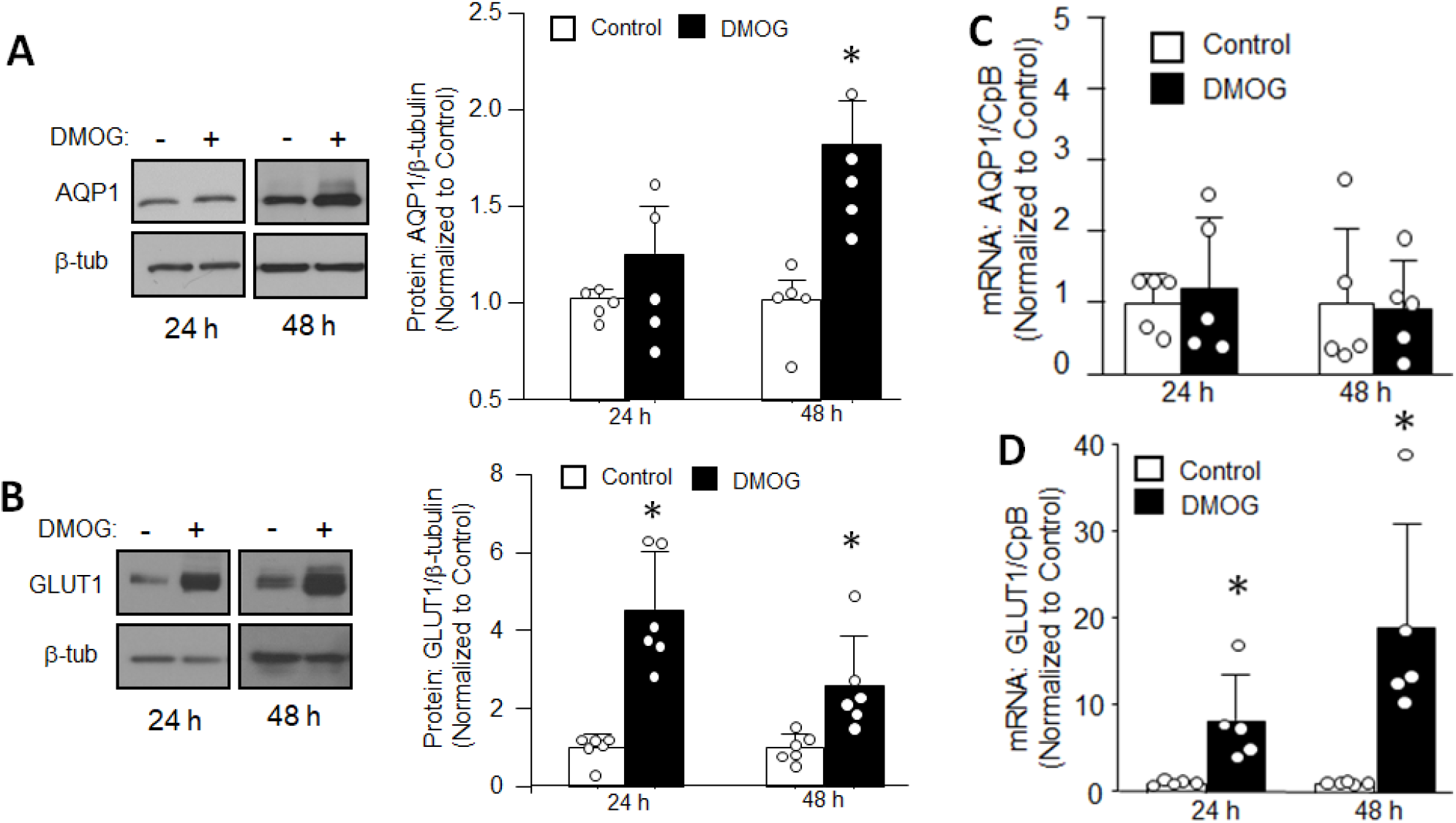
Effect of increasing hypoxia-inducible factors (HIFs) on aquaporin 1 (AQP1) and glucose transporter 1 (GLUT1) expression in pulmonary arterial smooth muscle cells (PASMCs). Representative immunoblot and bar (mean ± SD) and scatter plots showing A) AQP1 and B) GLUT1 protein levels in control cells treated with vehicle (DMSO; 1:3300) or DMOG (1 mM) for 24 and 48 h. * indicates p=<0.05 from Control at same time point by t-test. C and D) Bar (mean ± SD) and scatter plots showing AQP1 and GLUT1 mRNA expression normalized to cyclophīlin B (CpB) in PASMCs from control rats treated with vehicle or DMOG for 24 and 48 h. * indicates p<0.01 from Control at same time point by t-test. Each dot represents a biological replicate using cells isolated from a different animal.

To confirm that the effect of DMOG on AQP1 protein levels was mediated by HIFs and to further clarify the specific roles of HIF-1 or HIF-2, we repeated DMOG exposure in the presence of HIF inhibitors. We found that inhibition of HIF-1 (CAY10585; 30 μM) or HIF-2 (TC-S 7009; 10 μM) had no effect on AQP1 protein levels in cells exposed to vehicle (**Fig 3**). However, inhibiting either HIF-1 or HIF-2 attenuated the DMOG-induced increase in AQP1 protein levels, with no change in AQP1 protein evident with combined HIF-1 and HIF-2 blockade.

**Figure 3.**
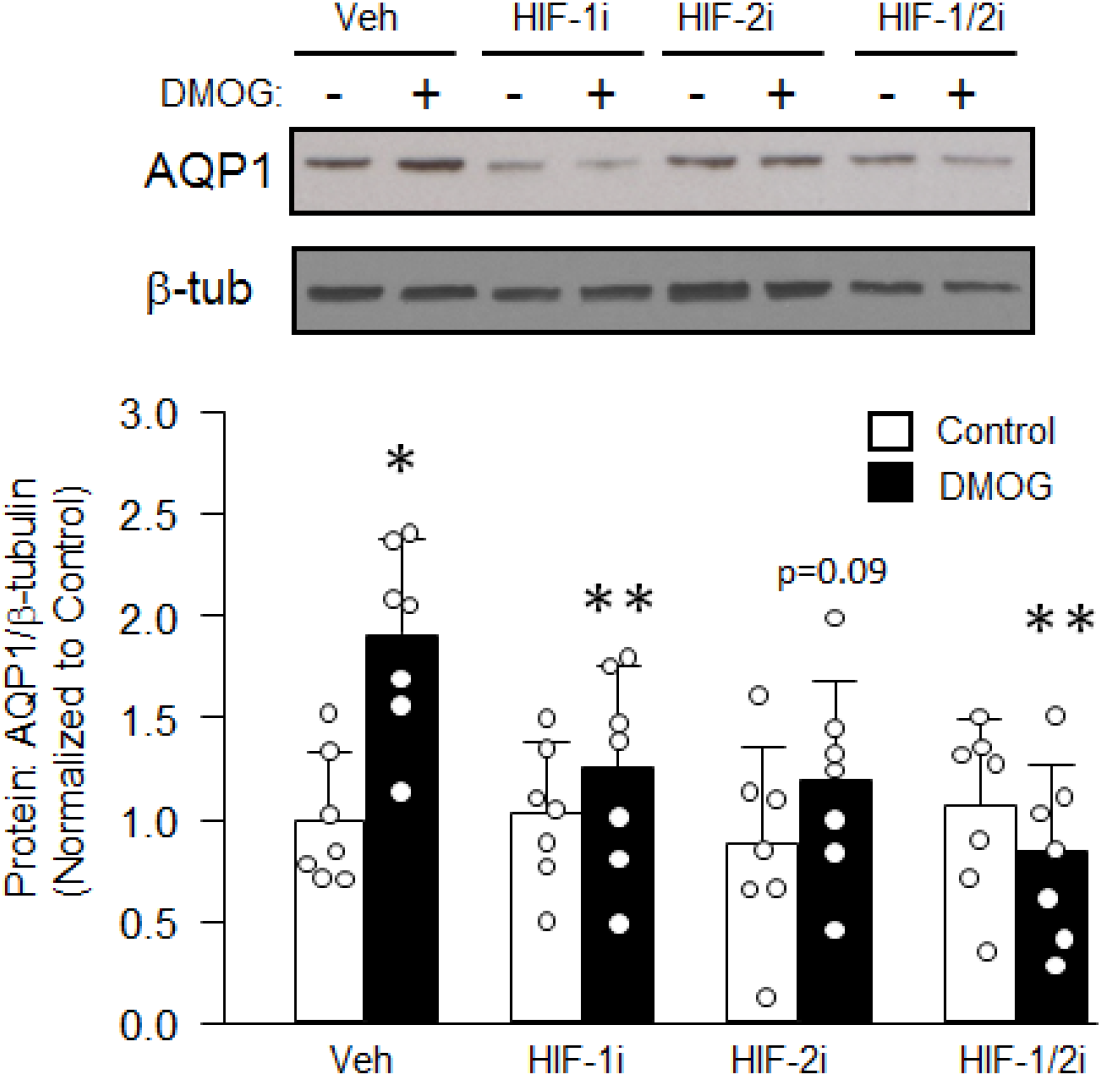
Effect of inhibiting hypoxia-inducible factors (HIFs) on DMOG-induced upregulation of aquaporin 1 (AQP1) protein expression in pulmonary arterial smooth muscle cells (PASMCs). Representative immunoblot and bar (mean ± SD) and scatter plots showing AQP1 and β-tubulin (β-tub; loading control) protein levels in control cells treated with vehicle (Control; 1:3300 DMSO) or DMOG (1 mM) for 48 h in the presence of vehicle (Veh; 1:2000 DMSO) or inhibitors of HIF-1 (HIF-1 i; 30 μM CAY10585), HIF-2 (HIF-2i; 10 μM TC-S 7009) or both. * indicates p=<0.05 from Control Veh; ** indicates p<0.05 from DMOG Veh by one-way ANOVA with Holm-Sidak post-test. Each dot represents a biological replicate using cells isolated from a different animal.

Because DMOG does not discriminate between HIF-1 and HIF-2, and HIF activation may not be its only action, we performed complimentary experiments using adenoviral vectors with constitutively active forms of HIF-1α (AdHIF-1α) and HIF-2α (AdHIF-2α). We measured AQP1 protein levels in cells infected with AdHIF-1α or AdHIF-2α, or a control construct containing EGFP (AdEGFP) for 48 h (**Fig 4A**). Augmenting HIF-1α or HIF-2α protein levels in the absence of hypoxia or DMOG was sufficient to increase mRNA levels of the canonical HIF targets GLUT1 and VEGF (**Fig 4B and C**). In contrast, AQP1 mRNA levels were unaltered by augmenting HIF-1α or HIF-2α. These results are similar to those observed with DMOG and provide further evidence that AQP1 is not a direct HIF transcriptional target in PASMCs. Despite the lack of change in AQP1 mRNA levels in cells with forced expression of HIF-1α or HIF-2α, AQP1 protein levels were increased (**Fig 4D**).

**Figure 4.**
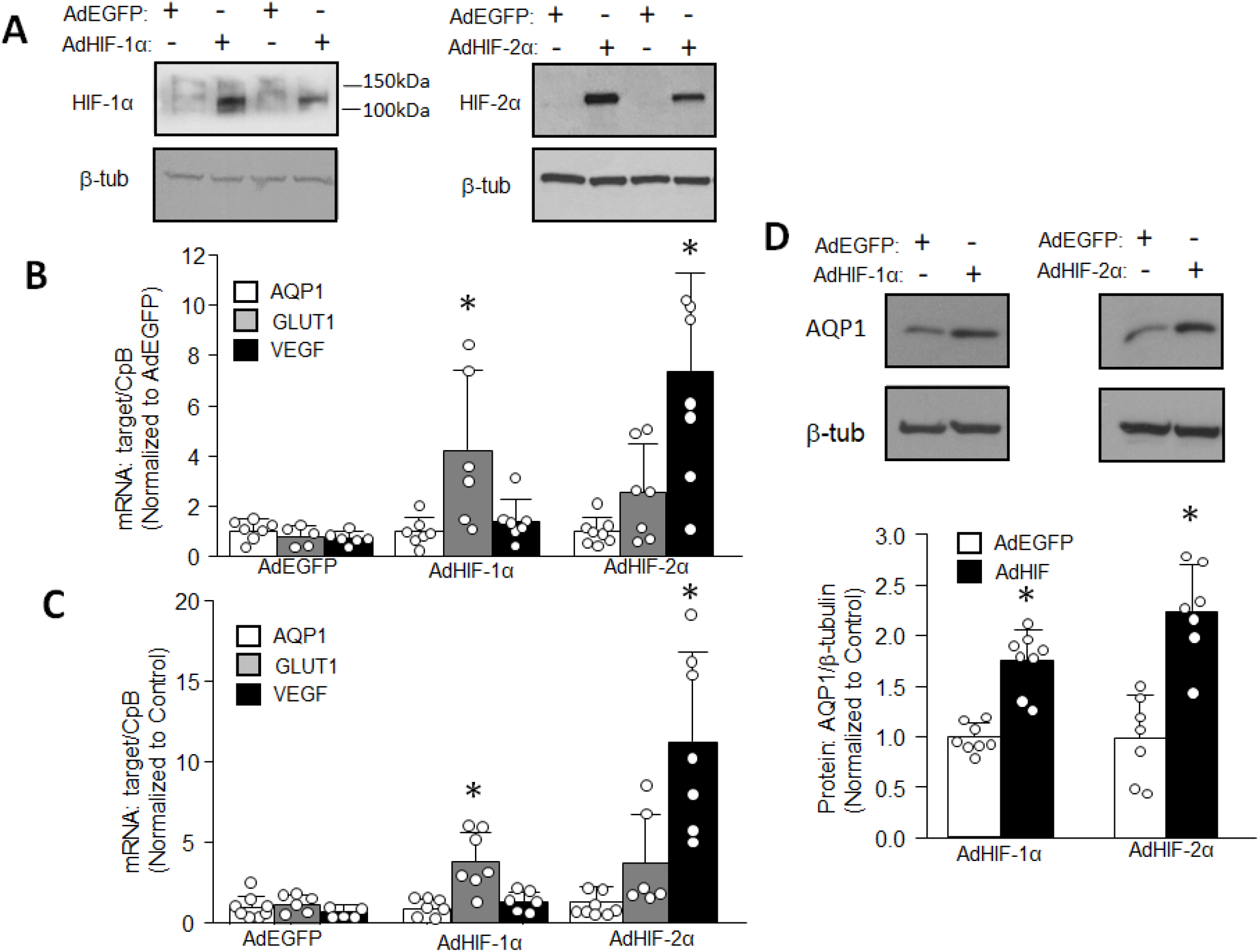
Effect of increased hypoxia-inducible factor (HIF) expression on pulmonary arterial smooth muscle cells (PASMCs). A) Representative immunoblots showing HIF-1α, HIF-2α and β-tubulin (β-tub; loading control) protein levels in control cells infected with AdEGFP (control) or adenovirus containing constitutively active HIF-1α (AdHIF-1α) or HIF-2α (AdHIF-2α) for 48 h. Bar (mean ± SD) and scatter plots showing mRNA levels for AQP1, the common HIF target GLUT1 and the primarily HIF-2 target, VEGF normalized to cyclophilin B (CpB) at B) 24 and C) 48 h after infection with AdEGFP, AdHIF-1α or Ad HIF-2α. * indicates p<0.05 from AdEGFP by oneway ANOVA. D) Representative immunoblots and bar (mean ± SD) and scatter plots showing AQP1 and β-tub protein levels in cells infected with AdEGFP, AdHIF-1α or AdHIF-2α for 48 h. * indicates p=<0.05 from AdEGFP by t-test. Each dot represents a biological replicate using cells isolated from a different animal.

### Effect of increasing HIF activity on intracellular Ca^2+^

Since the effect of DMOG on AQP1 expression did not appear to occur via direct transcriptional activation, we next explored potential intermediates that control AQP1 protein levels. We previously demonstrated that the effect of subacute hypoxia on AQP1 levels required an elevation in intracellular Ca^2+^ (16). We have also shown in PASMCs that the hypoxia-induced increase in [Ca^2+^]_i_ results from increased expression of Ca^2+^-permeable nonselective cation channels composed of TRPC proteins, the expression of which are under the control of HIFs (37). Thus, to determine whether an increase in [Ca^2+^]_i_ downstream of HIF activation was responsible for the augmentation in AQP1 protein in response to DMOG, we measured [Ca^2+^]i in PASMCs treated with DMOG or vehicle. We found that DMOG exposure for 48 h increased [Ca^2+^]_i_ (**Fig 5A**). HIF-2 inhibition partially reduced the DMOG-induced increase in [Ca^2+^]_i_, whereas the change in [Ca^2+^]_i_ in response to DMOG was completely prevented in the presence of the HIF-1 inhibitor alone or in combination with the HIF-2 inhibitor.

**Figure 5.**
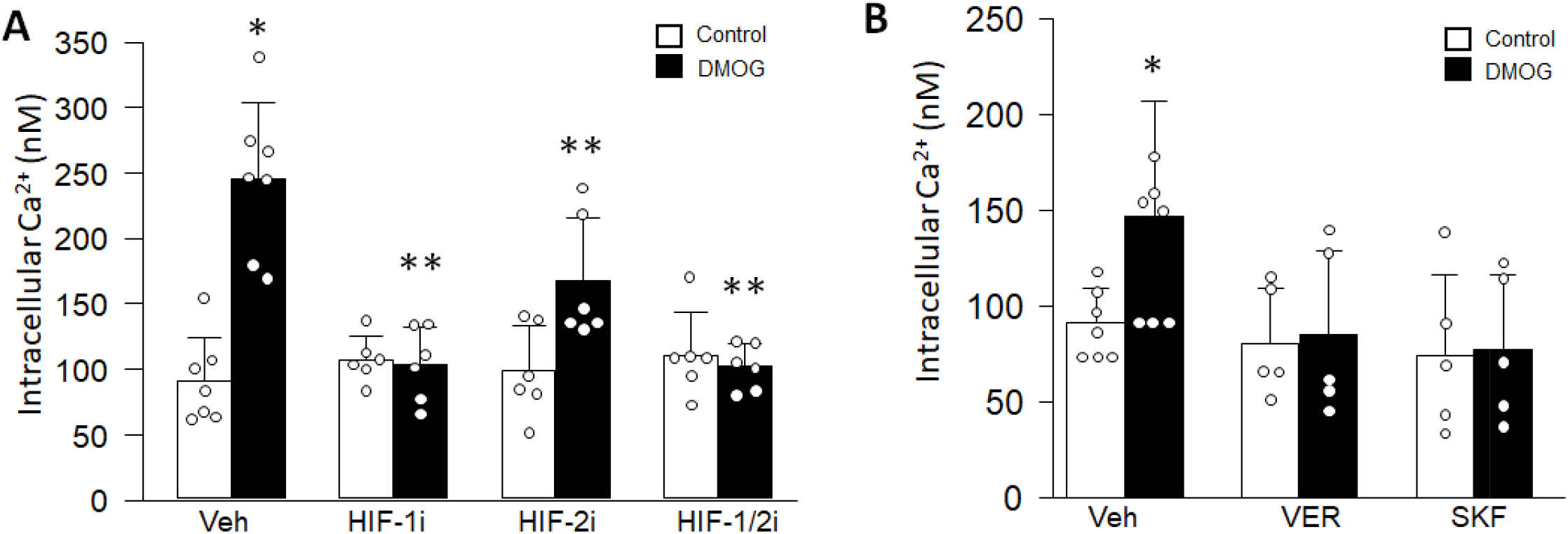
Effect of DMOG on intracellular Ca^2+^ concentration in pulmonary arterial smooth muscle cells (PASMCs). A) Bar (mean ± SD) and scatter plots showing baseline intracellular Ca^2÷^ concentration in PASMCs treated with vehicle (Control; 1:3300 DMSO) or DMOG (1 mM) for 48 h in the absence (Veh; 1:2000 DMSO) or presence of HIF-1 inhibitor (HIF-1 i; 30 μM CAY10585), HIF-2 inhibitor (HIF-2i; 10 μM TC-S 7009) or both. * indicated p<0.05 from Control Veh; ** indicates p<0.05 from DMOG Veh via ANOVA with Holm-Sidak post-test B) Bar (mean ± SD) and scatter plots showing baseline intracellular Ca^2+^ concentration in PASMCs treated with vehicle (Control) or DMOG for 48 h in the presence of (Veh; 1:2000 DMSO), the L-type Ca^2+^ channel blocker, verapamil (VER; 10 μM), or the nonselective cation channel blocker, SKF96365 (SKF; 30 μM). * indicates p<0.05 from Control under the same conditions by t-test. Each dot represents a biological replicate using cells isolated from a different animal.

We next identified the Ca^2+^ signaling pathways responsible for the DMOG-induced increase in [Ca^2+^]_i_. Similar to results we observed with moderate hypoxia in PASMCs (16, 36), both the L-type Ca^2+^ channel inhibitor, verapamil (VER; 10 μM) and the nonselective cation channel inhibitor, SKF96365 (SKF; 30 μM), prevented the change in [Ca^2+^]_i_ in response to DMOG, suggesting that Ca^2+^ influx through both channels can mediate the response (**Fig 5B**).

### Role of Ca^2+^ influx in modulating HIF-dependent increases in AQP1 protein levels

Since blockade of either L-type or nonselective cation channels prevented the increase in [Ca^2+^]_i_ in response to DMOG, our next step was to determine whether inhibiting these channels would also prevent the effects on AQP1 protein levels. While DMOG treatment resulted in a two-fold increase in AQP1 expression in vehicle-treated cells, no change in AQP1 protein was observed in response to DMOG in the presence of either VER or SKF (**Fig 6A**). In contrast, the increase in GLUT1 protein levels induced by DMOG was not altered by either Ca^2+^ channel blocker (**Fig 6B**). These data suggest that DMOG increases AQP1 protein levels via a mechanism involving increases in [Ca^2+^]_i_ driven by influx through L-type channels and NSCCs.

**Figure 6.**
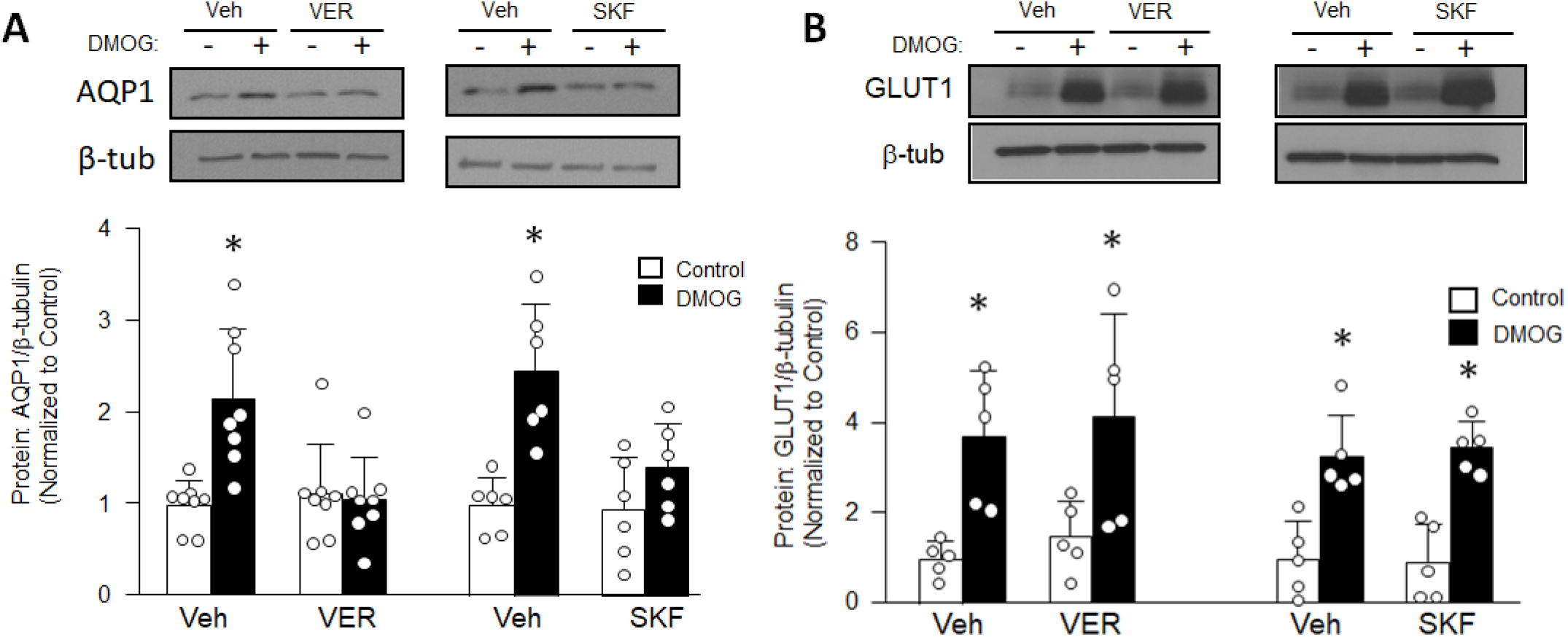
Effect of Ca^2+^ channel inhibitors on DMOG-induced increases in AQP1 and GLUT 1 protein expression in pulmonary arterial smooth muscle cells (PASMCs). Representative immunoblots and bar (mean ± SD) and scatter plots showing A) AQP1 and β-tubulin (β-tub; loading control) protein levels and B) GLUT1 and β-tub protein levels in cells treated with vehicle (Control: 1:33OO DMSO) or DMOG (1 mM) in the presence of vehicle (Veh: 1:2000 DMSO), the L-type Ca^2+^ channel inhibitor, verapamil (VER; 10 μM), or the nonselective cation channel inhibitor, SKF96365 (SKF; 30 μM) for 48 h. * indicates p<0.05 from Control under the same condition via 2way ANOVA with Holm-Sidak post-test. Each dot represents a biological replicate using cells isolated from a different animal.

### HIF1/2 gain-of-function and [Ca^2+^]_i_

To corroborate the findings with DMOG, we tested whether forced expression of HIF-1 and/or HIF-2 was sufficient to increase [Ca^2+^]_i_ in our cells and mimic the effects of DMOG. Compared to PASMCs infected with EGFP, infection with either AdHIF-1α (**Fig 7A**) or AdHIF-2α (**Fig 7B**) elevated basal [Ca^2+^]_i_ to levels similar to those observed with hypoxia (16, 37) or DMOG (**Fig 5**). Moreover, the increases in [Ca^2+^]_i_ induced by forced expression of HIF-1/2 could be prevented by treating the cells with VER or SKF, similar to the results observed in DMOG-treated cells. Consistent with the effects on [Ca^2+^]_i_, the increase in AQP1 protein levels induced by AdHIF-1α and AdHIF-2α were also prevented in the presence of VER or SKF (**Fig 8A and B**), indicating that the effects of HIF on AQP1 protein expression are mediated via increases in [Ca^2+^]_i_.

**Figure 7.**
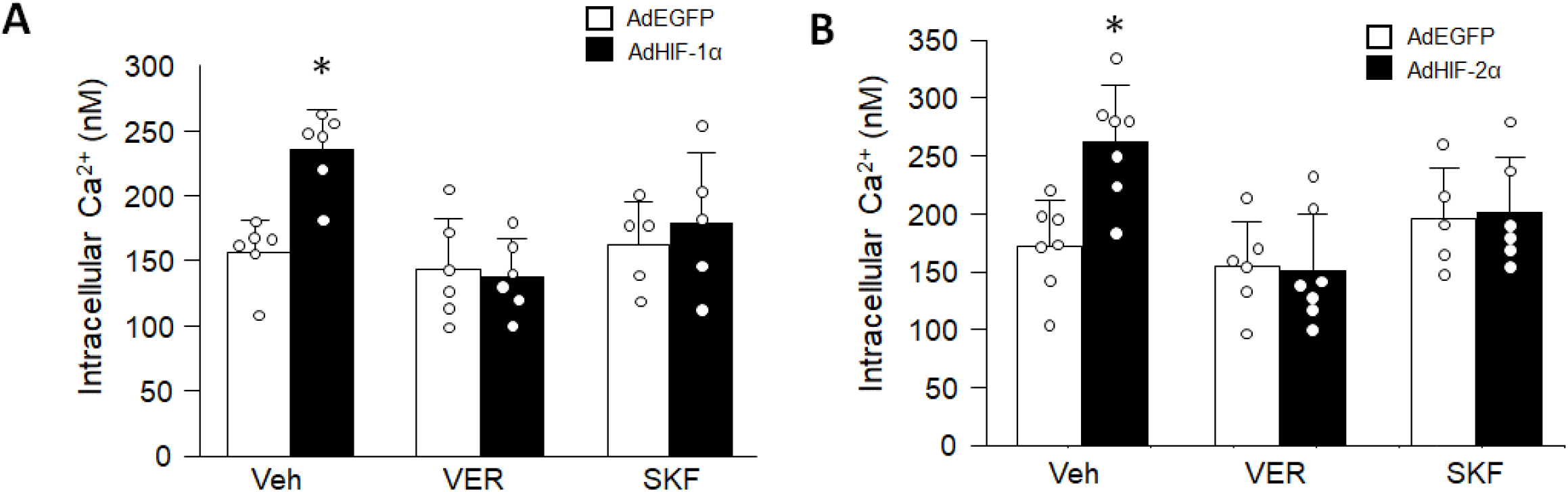
Effect of increasing HIF-1 and HIF-2 on intracellular Ca^2+^ concentration in pulmonary arterial smooth muscle cells (PASMCs). Bar (mean ± SD) and scatter plots showing intracellular Ca^2+^ concentration in PASMCs infected with the control adenovirus (AdEGFP) or adenovirus containing constitutively active A) HIF-1α (AdHIF-1α) or B) HIF-2α (AdHIF-2α)for 48 h in the presence of (Veh; 1:2000 DMSO), the L-type Ca^2+^ channel blocker, verapamil (VER; 10 μM), or the nonselective cation channel blocker, SKF96365 (SKF; 30 μM). * indicates p<0.05 from AdEGFP under the same condition via ANOVA with Holm-Sidak post-test. Each dot represents a biological replicate using cells isolated from a different animal.

**Figure 8.**
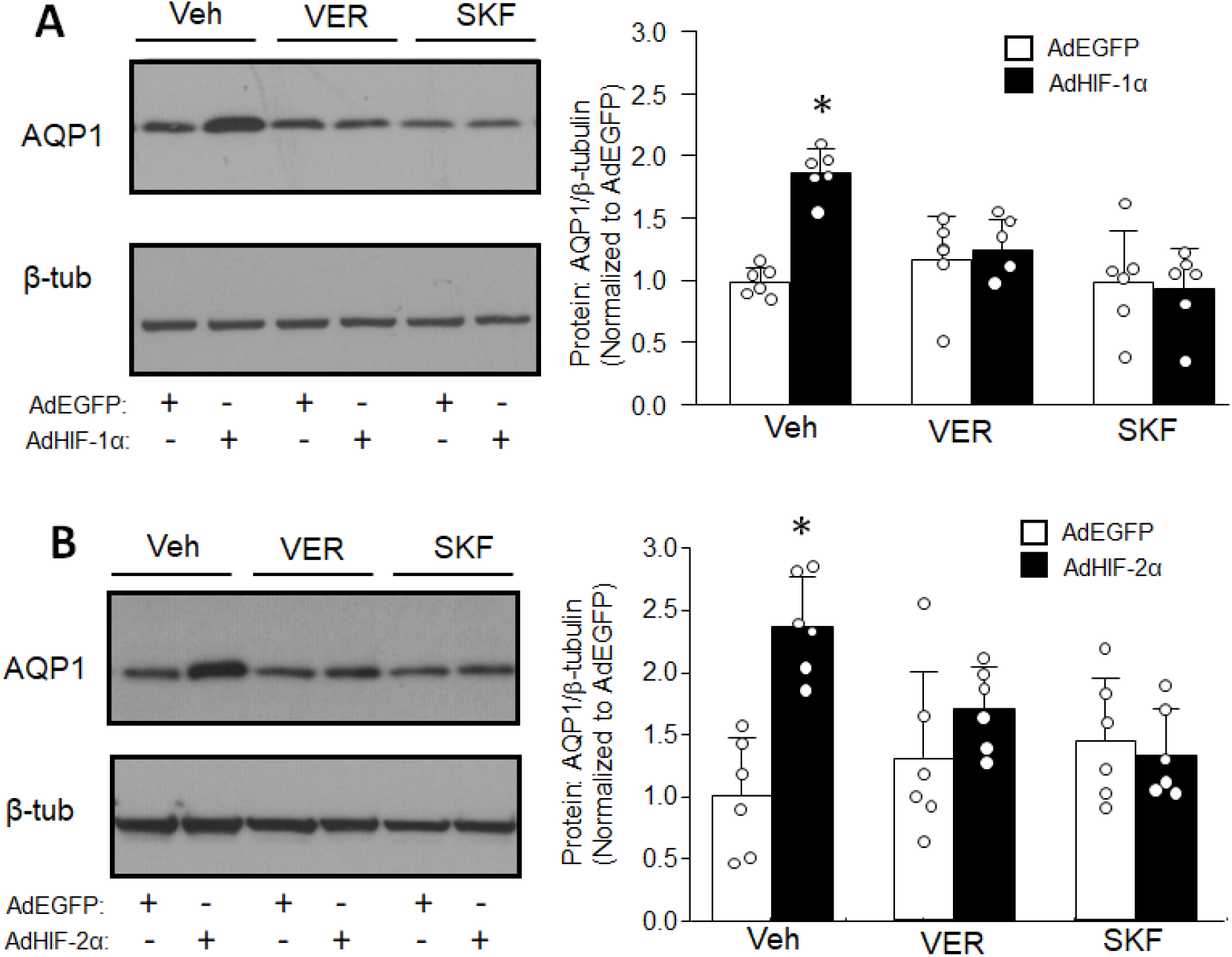
Effect of Ca^2+^ channel inhibitors on HIF-induced increases in AQP1 protein expression in pulmonary arterial smooth muscle cells (PASMCs). Representative immunoblots and bar (mean ± SD) and scatter plots showing AQP1 and β-tubulin (β-tub; loading control) protein levels in cells infected with AdEGFP or adenovirus containing constitutively active A) HIF-1α (AdHIF-1α) or B) HIF-2α (AdHIF-2α) for 48 h in the presence of vehicle (Veh: 1:2000 DMSO), the L-type Ca^2+^ channel blocker, verapamil (VER; 10 μM), or the nonselective cation channel blocker, SKF96365 (SKF; 30 μM). * indicates p<0.05 from AdEGFP under the same condition via ANOVA with Holm-Sidak post-test. Each dot represents a biological replicate using cells isolated from a different animal.

### Increased HIF1/2 and PASMC function

Finally, we tested whether activation of HIF-1 or HIF-2 was sufficient to increase PASMC migration and proliferation, and if so, whether this required AQP1. In PASMCs where HIF-1 or HIF-2 was increased under non-hypoxic conditions via AdHIF-1α and AdHIF-2α, migration and proliferation were increased compared to PASMCs infected with AdEGFP (**Fig 9**). After depleting AQP1 with siRNA, AdHIF-1α and AdHIF-2α no longer increased migration or proliferation. These results indicate that increasing HIFs in the absence of hypoxia is sufficient to induce changes in PASMCs function via mechanisms requiring increased AQP1.

**Figure 9.**
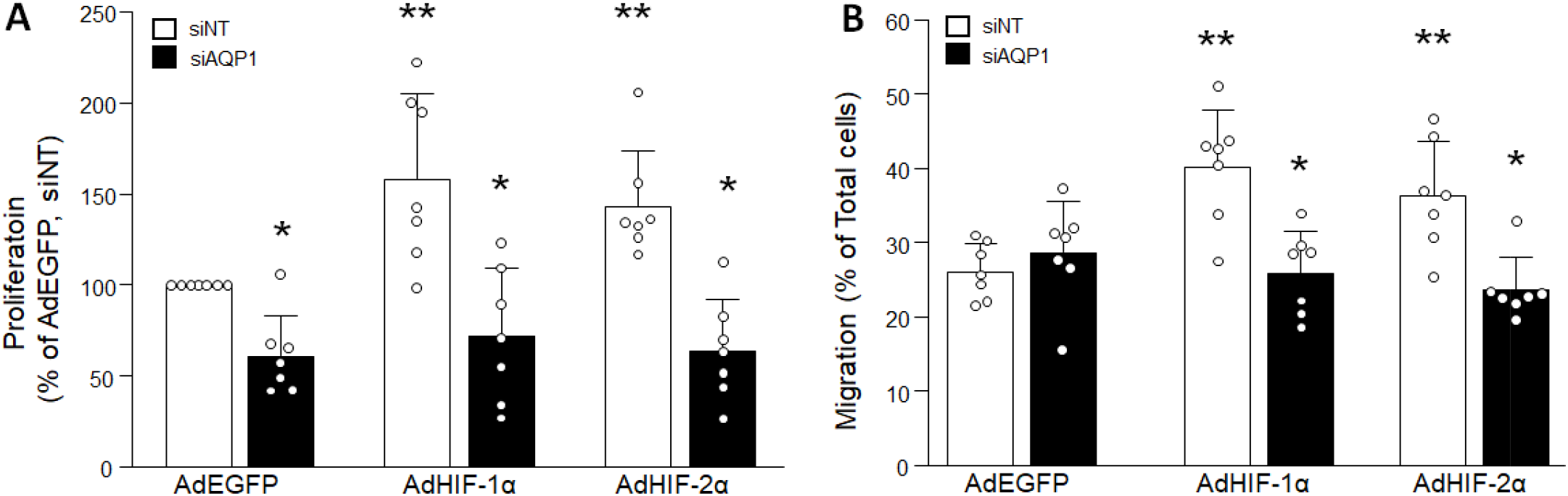
Effect of silencing AQP1 on HIF-induced increases in proliferation and migration in pulmonary arterial smooth muscle cells (PASMCs). A) Bar (mean ± SD) and scatter plots showing proliferation measured in PASMCs infected with adenovirus containing EGFP (AdEGFP; control) or constitutively active HIF-1 or (AdHIF-1α) or HI F-2α (AdHIF-2α) when cells were transfected with nontargeting siRNA (siNT) or siRNA targeting AQP1 (siAQP1). B) Bar (mean ±SD) and scatter plots showing migration measured in PASMCs infected with adenovirus containing EGFP (AdEGFP; control) or constitutively active HIF-lα (AdHIF-1α) or HIF-2α (AdHIF-2α) in cells were transfected with nontargeting siRNA ¢siNT) or siRNA targeting AQP1 (sīAQPI). In both graphs, * indicates p<0.05 from sīNT under the same condition and “ indicates p<0.05 from AdEGFP/sīNT group via 2-way ANOVA with Holm-Sidak post-test. Each dot represents a biological replicate using cells isolated from a different animal.

## DISCUSSION

AQP1 plays a role in several vascular functions, including regulation of migration and proliferation of both PASMCs (15, 16, 18, 25, 40) and ECs (23, 34). We have previously demonstrated that AQP1 was required for the hypermigratory and hyperproliferative PASMC phenotype induced by hypoxia (16), a finding subsequently corroborated by others (18, 25). HIF-1 has also been implicated in mediating hypoxia-induced migration and proliferation in PASMCs (27). Given the similar roles these proteins play in regulating cell phenotype and previous work in retinal endothelial (31) and hemangioendothelioma (1) indicating a functional HIF binding site in the AQP1 promoter it seemed reasonable that HIF may directly regulate AQP1 transcription in PASMCs. In this study, we used a variety of approaches to activate HIFs, including exposure to hypoxia, inhibiting prolyl hydroxylases with DMOG and forced expression of HIFα proteins, and measured the effect on AQP1 mRNA and protein levels. In contrast to results obtained in other cell types, we found that activation of either HIF-1 or HIF-2 increased AQP1 protein but had no effect on AQP1 mRNA. The augmentation of AQP1 protein abundance appears to be mediated by Ca^2+^-dependent post-transcriptional mechanisms involving both L-type and nonselective cation channels rather than HIF-dependent transcriptional regulation.

Taking advantage of tissues banked from previous experiments, we examined the effect of treatment with digoxin, which inhibits HIF-1α accumulation (42), on chronic hypoxia-induced AQP1 protein expression in lung tissues. Consistent with our previously reported results demonstrating treatment with digoxin prevented hypoxia-induced upregulation of other HIF targets in these tissues (2), the hypoxia-induced increase in AQP1 protein was also prevented by digoxin-treatment. These experiments are limited in that use of whole lung tissue does not allow identification of the specific cell type involved and we cannot rule out off-target effects of digoxin. Also, since small amounts of tissue were available, we were unable to also measure mRNA levels. Thus, it remains unknown whether AQP1 mRNA levels were increased by chronic hypoxia in these particular tissues, as has been reported previously for hypoxic rats and mice (1, 9, 16), or whether digoxin treatment transcriptionally inhibited AQP1. Nonetheless, these experiments confirm previous findings of increased AQP1 protein levels in the lungs of hypoxic mice (18, 25) and rats (16) and provide proof-of-concept that elevations in lung AQP1 levels during in vivo CH could involve HIF.

While PASMCs certainly sense and respond to hypoxia with an increase in HIF accumulation (24), whether the upregulation of AQP1 protein observed in PASMCs from chronically hypoxia animals was HIF-1 dependent was unknown. To more directly assess the role of HIFs in regulating AQP1 expression in PASMCs, we used cells exposed to ex vivo hypoxia, a prolyl hydroxylase inhibitor or infected with viral constructs that increased HIFα levels. In all cases, protein levels of both AQP1 and the canonical HIF target, GLUT1, were increased, confirming that HIFs modulate AQP1 protein expression. However, while GLUT1 mRNA levels were robustly increased within 24 h, there was no consistent effect on AQP1 mRNA levels. Although these results are in line with previous studies demonstrating modest (1) or variable (25) effects of hypoxia on AQP1 mRNA in cultured PASMCs, it was nonetheless a surprising finding given that previous studies indicated that AQP1 was a direct HIF target in certain cell types (1, 31). Moreover, we previously demonstrated that exposing cells from normoxic animals to hypoxia ex vivo increased AQP1 protein, but not mRNA levels (16). This intriguing dichotomy between the effects of in vivo and in vitro hypoxia exposure raised the possibility that AQP1 might not be a direct HIF target gene in all cell types. Use of PASMCs from control animals where we artificially increased expression of HIF-1α and HIF-2α, either by inhibiting PHD activity or expressing HIF-1α and/or HIF-2α proteins that were unable to be degraded, allowed us to directly assess the effects of HIF in the absence of hypoxia. Interestingly, we found no change in AQP1 mRNA or protein levels at 24 h, compared to the robust upregulation observed for the canonical HIF target, GLUT1. That GLUT1 levels increased as expected suggests that HIF activation occurred as anticipated, but simply failed to increase AQP1 levels at early timepoints. This finding is consistent with the lack of early effects of hypoxia on AQP1 protein levels in human PASMCs exposed to hypoxia in vitro, despite substantial accumulation of HIF-1α protein at much earlier timepoints (25). We observed reliable increases in AQP1 protein levels at 48 h, but again with no detectable corresponding change in mRNA levels. These results with HIF activation are consistent with the previously reported effects of ex vivo subacute exposure to moderate hypoxia in rat PASMCs (16), which increased AQP1 protein expression without a concomitant increase in mRNA levels.

What remains unclear is whether the increase in AQP1 mRNA observed in PASMCs from chronically hypoxic rats results from increased HIF activation or occurs via another mechanism. Given that directly increasing either HIF-1 or HIF-2 had no effect on AQP1 mRNA, we suspect that the latter is the more plausible possibility, although further experiments will be required to definitively answer this question. Also remaining to be determined is whether the increase in AQP1 mRNA observed with in vivo hypoxic exposure (1, 16) was due to the extended duration of hypoxic exposure or to other factors only found in vivo (i.e., circulating or paracrine factors, mechanical stress, interactions with the matrix, etc). Future experiments will be needed to determine the exact factors involved in, and timing of, the transcriptional upregulation of APQ1 in chronically hypoxic animals. Nonetheless, when combined with the HIF-dependent increase in AQP1 protein accumulation, an additional increase in mRNA would provide a mechanism for robust upregulation of AQP1 during the development of hypoxic pulmonary hypertension.

Our findings indicate that both HIF-1 and HIF-2 can be involved in the hypoxia-induced increase in AQP1 protein, as inhibiting either was sufficient to prevent changes in AQP1 abundance with hypoxia. Similarly, increasing either HIF-1 or HIF-2 enhanced AQP1 protein levels. In our in vivo studies, at the concentrations used, digoxin is believed to be more selective for HIF-1α (42), but we cannot rule out that HIF-2α was also inhibited in these studies. We also found a role for both HIF-1 and HIF-2 in regulating PASMC [Ca^2+^]_i_. Our loss-of-function experiments showed that inhibiting either HIF-1 or HIF-2 was sufficient to normalize [Ca^2+^]_i_ in the presence of DMOG. Interestingly, in our previous experiments, treatment with digoxin, which reduced lung AQP1 levels, also normalized [Ca^2+^]_i_ in PASMCs from chronically hypoxic mice (2). On the other hand, gain-of-function of either HIF-1α or HIF-2α increased PASMC [Ca^2+^]_i_ levels to values similar to those observed with DMOG. These results are consistent with our previous work showing that HIF-1 could upregulate canonical transient receptor potential (TRPC) proteins (37), which form nonselective cation channels (NSCCs), providing a mechanism by which DMOG could regulate Ca^2+^ influx. Indeed, we found the increase in [Ca^2+^]_i_ induced by DMOG was prevented when cells were treated with an inhibitor of NSCCs. Interestingly, inhibition of voltage-gated Ca^2+^ channels (VDCC) also normalized [Ca^2+^]_i_ in DMOG-treated cells and in cells where HIF-1α or HIF-2α was increased. Earlier data demonstrated that acute hypoxia increases the activity both channel types (36), while subacute exposure to hypoxia in vitro upregulates the expression of TRCP1 and TRPC6 in a HIF-dependent manne r(37). One explanation could be that activation of TRPC channels, while allowing influx of Ca^2+^ directly, could also result in depolarization due to changes in Na^+^ and/or K^+^ concentrations, which in turn could activate VDCC. It is also possible that HIF activation downregulated K^+^ channel activity and/or expression, as we (29, 38) and others (5) have previously demonstrated. A reduction in K^+^ channel activity would lead to depolarization of PASMCs, and influx through VDCC. However, inhibitors of both VDCCs and NSCCs were equally effective in preventing the HIF-induced increase in [Ca^2+^]_i_, implying that the role of K^+^ channels in activating VDCCs independent of NSCCs might be minimal in our cells.

It should be noted that in our previous work (37), we were unable to test whether forced expression of HIF-1 increased [Ca^2+^]_i_ since our control virus included GFP, which interferes with Fura-2 measurements due to overlapping excitation/emission spectra. In the current study, we purposely chose EGFP as our control since the wavelengths for excitation do not overlap with Fura-2. While the increase in [Ca^2+^]_i_ induced by HIF activation observed in this study is consistent with our previous findings showing loss of HIF-1α reduced PASMC [Ca^2+^]_i_ during hypoxia (37), these results contrast with other reports in human PASMCs showing that loss of HIF increased hypoxia-induced elevations in [Ca^2+^]_i_ (4). The reasons for this discrepancy are unclear, but could relate to a variety of factors, including sex of the subjects, species, culture conditions and/or location within the vasculature from which the cells were derived.

The exact mechanism by which increased expression of HIFs and elevated [Ca^2+^]_i_ increased AQP1 protein levels in our cells remains to be determined. We attempted to determine whether the increase in AQP1 protein was a result of increased protein stability (i.e., reduced degradation); however, we found that the half-life of the protein in our cells was >24 h (data not shown) and that exposure to cycloheximide for this duration caused cell death and detachment. The half-life of the AQP1 protein in PASMCs would appear to be much longer than has been reported in other cell types (17, 33) although the reason for this apparent discrepancy is unclear. Another potential point of interest is that while the current study explored regulation of AQP1 protein abundance by HIFs, a previous study demonstrated hypoxia-induced increases in HIF-1α expression were prevented when AQP1 was depleted (18). We did not test the effect of altering AQP1 levels on HIF expression in our study, but in total, these findings could suggest a potential feed-forward mechanism, whereby hypoxia may serve as an inciting factor to induce HIFs, with the subsequent accumulation of AQP1 further augmenting HIF activation, providing a mechanism to maintain HIF activation even in the absence of continuing hypoxia.

We previously reported that exposure to hypoxia increased PASMC proliferation and migration (35), contributing to vascular remodeling during pulmonary hypertension. Studies have shown that the effect of hypoxia on PASMC proliferation is HIF-(12, 27) and Ca^2+^-dependent (20). We and others have also demonstrated a necessary role for AQP1 in hypoxia-induced changes in PASMC proliferation. Thus, to assess the functional role of HIF-induced AQP1 protein accumulation we tested the effect of directly increasing HIFs in the absence and presence of AQP1. We found that enhancing HIF expression was sufficient to induce a pro-migratory, pro-proliferative phenotype in PASMCs, and that depleting AQP1 could prevent these changes in cell phenotype. These results likely explain why both loss of HIF-1 in SMCs and loss of AQP1 similarly reduced remodeling and development of PH in animal models (3, 18).

A limitation of the current study is that our work was performed primarily in culture models and may not entirely reflect the in vivo conditions. However, given that AQP1 protein levels are increased in the lungs and cells (16) from hypoxic animals, that loss of AQP1 reduced hypoxia-induced PH (18, 25), and that inhibition or deletion of HIF-1 or HIF-2 in vivo is most often shown to be protective against the development of PH (3, 6, 8, 13, 14, 39), our results are consistent with the possibility that HIF-dependent upregulation of AQP1 in PASMCs could be contributing to the vascular remodeling observed in vivo.

In summary, we describe the induction of AQP1 protein accumulation by both HIF-1 and HIF-2 in PASMCs. Despite previously identified HIF binding sites in the *Aqp1* promoter, in PASMCs it appears that HIFs do not exert direct transcriptional regulation of AQP1. Building on studies showing HIF1/2 depletion or deletion in mice results in reduced PH in response to hypoxia (3, 6, 13, 14, 39), interest in HIF inhibitors as therapeutic options for pulmonary hypertension have garnered substantial recent interest (2, 8, 13, 19). A limitation of the current study is that although it is clear that HIFs can regulate AQP1 protein expression in PASMCs, which HIF is acting in vivo cannot be determined from these studies. Additional work measuring AQP1 expression in PASMCs in animals treated with specific HIF-1 or HIF-2 inhibitors will be required. However, the results of the current study would seem to suggest a component of the benefit offered by HIF inhibition in pre-clinical models could stem from effects on AQP1 protein levels. While chronic hypoxia was used as stimulus and starting point for the current studies, studies have reported increased HIF activation/expression has been described in other types of PH where hypoxia is not a contributing factor, such as pulmonary arterial hypertension (10, 32), suggesting the potential for AQP1 to play a role in other forms of the syndrome. Moreover, an AQP1 variant was found to be associated with development of PAH (11), although whether this variant confers a gain-of-function mutation is at present unclear. Together, these findings would suggest that AQP1 plays a critical role in various forms of PH, further pointing to the need to understand its role and regulation in PASMCs and other cell types in the quest for developing new therapeutics that could target vascular remodeling.

## ACKNOWLEDGEMENTS

This work was funded by grants from the National Institutes of Health (R01HL073859, R01HL126514 and R25 HL084762) and the American Heart Association (18POST34030262).

## Notes

### Competing Interest Statement

The authors have declared no competing interest.

### Summary of Updates

Updated author list

